# Choice of method of place cell classification determines the population of cells identified

**DOI:** 10.1101/2021.02.26.433025

**Authors:** D.M. Grijseels, K. Shaw, C. Barry, C.N. Hall

## Abstract

Place cells, spatially responsive hippocampal cells, provide the neural substrate supporting navigation and spatial memory. Historically most studies of these neurons have used electrophysiological recordings from implanted electrodes but optical methods, measuring intracellular calcium, are becoming increasingly common. Several methods have been proposed as a means to identify place cells based on their calcium activity but there is no common standard and it is unclear how reliable different approaches are. Here we tested three methods that have previously been applied to two-photon hippocampal imaging or electrophysiological data, using both model datasets and real imaging data. These methods use different parameters to identify place cells, including the peak activity in the place field, compared to other locations (the Peak method); the stability of cells’ activity over repeated traversals of an environment (Stability method); and a combination of these parameters with the size of the place field (Combination method). The three methods performed differently from each other on both model and real data. The Peak method showed high sensitivity and specificity for detecting model place cells and was the most robust to variations in place field width, reliability and field location. In real datasets, vastly different numbers of place cells were identified using the three methods, with little overlap between the populations identified as place cells. Therefore, choice of place cell detection method dramatically affects the number and properties of identified cells. We recommend the Peak method be used in future studies to identify place cell populations, unless there is an explicit theoretical reason for detecting cells with more narrowly defined properties.

**Author Summary:** Place cells are hippocampal cells that have spatially constrained receptive fields, the place field. These cells have been widely studied in the context of navigation, more recently using virtual reality environments in combination with optical methods of recording neuronal activity. However, there is a lack of consensus regarding how to identify place cells in these data. In this study we tested the sensitivity and specificity of three methods of identifying place cells. By comparing these methods and quantifying the populations of place cells they identify, we aimed to increase our understanding of exactly the populations that are currently being studied under the name “place cells”. Although the appropriate method may depend on the experimental design, we generally recommend a single method going forward, which will increase consensus within the field about what should be included in a place cell population, and allow us to better compare results between studies.

## Introduction

Place cells are a subset of hippocampal pyramidal cells that fire selectively when the subject is in a certain location [1], and provide a sparse population code for self-location. Studies revealing their properties, including location-specific firing [1], directional selectivity [2] and context-dependence [3] have been vital for our understanding of how the hippocampus codes space. [4]. Place cells are characterised by their place field, which is a spatially stable location where the cell preferentially fires. Depending on the size of the environment, place cells can have one or multiple place fields [5]. The place fields may change location between different environments, a phenomenon called remapping. Place cells are also relevant to understanding clinical conditions such as Alzheimer’s disease, as in mouse models of Alzheimer’s disease they show impaired firing [6] linked to memory deficits.

Since their discovery, place cells have been extensively studied in real world environments – both open field and constrained - using electrophysiological recordings while animals either explore freely or perform directed spatial tasks. In these studies, cells are generally included in analyses based on their peak firing rate and stability, and the place fields of these cells are identified based on their spatial properties (e.g. [7]). These place fields are often characterised on functional basis, such as by the peak firing rate occurring in a given location, in otherwise sparse firing [8], and the stability of this place field over time (see e.g. [9]).

However, methodological advances now also allow place cells to be studied in vivo using calcium imaging [10–12], enabling large populations (n>100) of cells to be recorded for multiple sessions. This method requires the brain to be stationary during recording, which necessitates the use of a Virtual Reality (VR) environment. Often a visual VR is used during imaging, consisting of one or multiple screens displaying an environment that the mouse can control by running on a ball or wheel. With a ball, the mouse is able to move in two dimensions (e.g. [13]), whereas with a wheel, mice can only move backwards and forwards. These types of VR limit the animal’s ability to look around, and provide sparse sensory feedback; for example, whisking does not provide information about the environment. Place cells also respond differently in VR environments compared to the real world, showing broader place fields and increased directionality [14].

Several studies using one-dimensional environments (i.e. corridors) have revealed that hippocampal pyramidal cells represent other features in addition to location, such as reward [15] and travelled distance [16]. It is unclear to what extent these features are coded by separate populations of cells in the hippocampus or cells that, under other circumstances such as a dedicated navigation task, might be recruited as place cells. To resolve this, it is important to be able to accurately define place cells within such one-dimensional environments, but currently different studies use widely varying methods making comparisons between studies problematic.

Varying definitions of place cells have been chosen to account for the constraints imposed by the imaging methodology. Unlike electrophysiological recordings, imaging detects changes in intracellular calcium levels rather than direct readouts of action or synaptic potentials. The non-linear relationship between calcium transients and spike rates makes it hard to accurately estimate spike rates from two-photon data [17], so adopting exactly the same methods as used in electrophysiological recordings is not possible. Instead, studies have tended to use adaptations of some, but not all, of the imaging equivalents of peak firing rate, sparseness of off-location firing, and place field stability to define a cell as a place cell: Dombeck et al. [10] categorised a cell as a place cell based on a combination of properties of that cell’s apparent place fields, including the size, peak calcium fluorescence and the ratio of the firing within and outside the field. This method, or variations of it, have subsequently been adopted in several studies (e.g. [18, 19]). Fournier et al. [20] proposed a statistical method that uses the peak activity in the rate map of a cell and compares it to the peak activity in shuffled versions of the cell. An alternative approach [21] detects place cells from the stability of their activity as a function of location, a method based on those used in electrophysiology experiments [3]. However, it is unclear what biases these different methods exhibit and to what extent their classification criteria are equivalent – in short, do they identify the same neurons as place cells?

In this paper we aim to address the lack of consensus on how to identify place cells in two-photon data in a one-dimensional VR environment. We compare the performance of two established methods for the identification of place cells in two-photon data – the methods used by Dombeck et al. [10] and O’Leary et al. [21]– as well as one method developed for use on electrophysiology data which has been adapted for use in imaging – described by Fournier et al. [20] . We applied them to a range of synthetic model cell populations to explore whether cells identified as place cells by each of these methods possess similar characteristics. We find that in a range of mock datasets the method developed by Fournier et al., which we call the Peak method, is best able to identify place cells, having a high sensitivity and specificity, and detects more cells in real datasets. As a result, we recommend this method for the identification of place cells in two-photon imaging data. Our data also show that choice of identification method is important, as the methods classify different, largely non-overlapping populations of cells as place cells

## Results

### Place cell detection by the Combination and Stability methods is sensitive to the properties of place fields and the number of times mice traverse the environment

We evaluated the performance and suitability of three different approaches for detecting place cells (S1 Fig):

1. The Peak method, which classifies cells as place cells based on the average rate of firing (approximated by fluorescence change in two-photon data) in one location being higher than in the rest of the environment [22];
2. The Stability method, which identifies place cells as those with stable firing patterns across locations over time [21];
3. The Combination method, which requires place cells to fire more across a contiguous stretch of the environment than at baseline and to fire in this location on at least 20% of traversals [10].

To characterise the three different approaches for detecting place cells we applied them to synthetic datasets consisting of 20 true place cells and 80 random cells, matching the prevalence of place cells found using patch-clamp recordings [23]. Mouse location for each dataset was generated using experimentally acquired locomotion time courses from mice running through a linear virtual reality environment (S2 Fig, 8 datasets: 4 mice, 2 sessions per mouse). The dataset contained a total of 184 traversals of the environment, with mice running 23 traversals per session on average. Model fluorescence maps were generated by applying the synthetic cell profiles to 50 randomly selected traversals (Fig 1A,B). Periods where the mouse was not running – defined as having a velocity below 10% of the maximum – were removed (Fig 1C,D).

**Fig 1.**
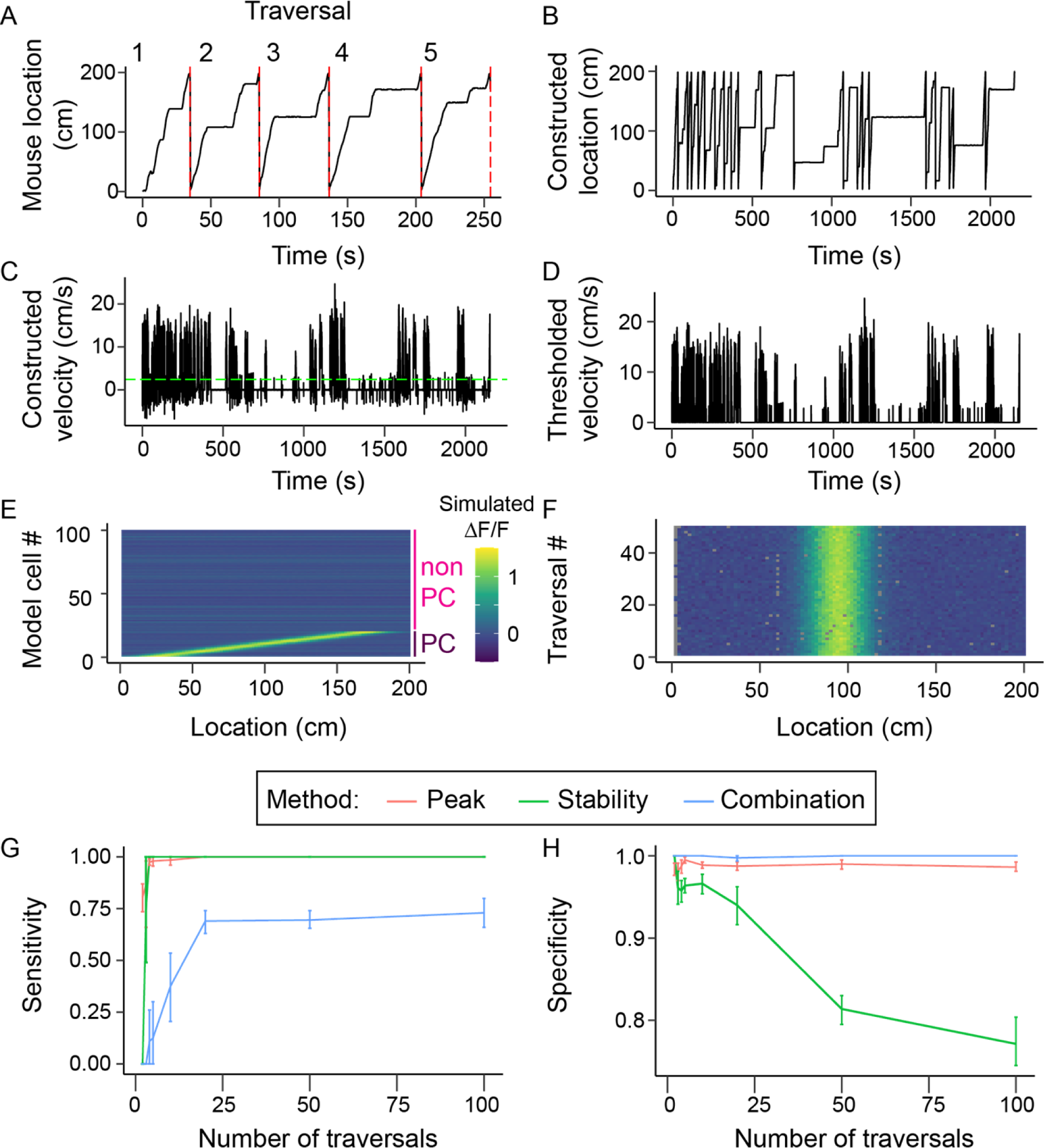
Model locomotion profile generation and effect on methods. (A) Section of a locomotion profile collected in a 200cm long virtual environment. This was cut into individual traversals of the environment (numbered, separated by red dashed lines) (B) locomotion profile generated from 20 randomly selected traversals. (C) Velocity profile of the locomotion profile shown in B. The green dashed line indicates the threshold for detecting running (at 10% of the maximum velocity) (D) Trace in C after thresholding. Time points with below threshold velocity were excluded from further analysis. (E) Fluorescence maps of 20 model place cells (cells 1-20) and 80 non-place cells (cells 81-100). (F) Activity over location of a single model place cell over 50 traversals. Grey points are missing data as the mouse moved across this location between acquisition of two frames. (G) The sensitivity of each method as a function of the number of traversals included in the locomotion trace. (H) The specificity of each method as a function of number of traversals. Lines in (G) and (H) show means over 10 randomly generated data sets. Error bars represent 95% confidence intervals.

Fluorescence maps of the place cells were modelled using a Gaussian to simulate changes in fluorescence as a function of location, where the mean determined the place field centre and sigma the field width. We used a default width of 50cm, based on widths previously reported in a virtual environments [10, 23]. Synthetic data was linearly scaled to match the peak values observed in our own two-photon CA1 data (Fig 1E). We distributed place field centres evenly across the 200 cm long track, then convolved the place field with the position of the mouse to determine the fluorescence map of each cell over time (Fig 1E, bottom 20 cells). We also included 80 control cells (Fig 1E, top 80 cells) with a fixed mean fluorescence but no place-dependence. Poisson was then added to all cells and the three different place cell classification methods were applied to these fluorescence maps.

We could therefore vary the parameters that determine the fluorescence map (Gaussian peak, width and location, number of traversals of the environment), to determine how much the properties of the place field affect their detection by the different methods. To assess place cell classification performance, we first calculated how many place and non-place cells were correctly identified by each method (true positives and true negatives), versus the number of false positive and false negative identifications. From these values, we then calculated the sensitivity (the proportion of place cells that were correctly detected), and the specificity (the proportion of cells identified as place cells that actually were place cells) of each method (Methods, equations (1) and (2)). A perfect detection method would therefore have a selectivity of 1 and a specificity of 1.

### Number of traversals

First, we determined the impact of the number of times the mouse ran through the environment on place cell detection by varying the number of traversals used to generate the model datasets from 2 to 100 (Fig 1G,H). We generated 10 model datasets for each number of traversals.

Both the Peak and the Stability method had a high sensitivity regardless of the number of traversals included in the dataset (Fig 1G). The Peak method also displayed a stable specificity across the range of traversals, with a mean of 0.99, reflecting its requirement that a true place cell’s activity is in the top 1% of shuffled data (Fig 1H). However, the Stability method saw a decreasing specificity with an increase in traversals, with a specificity of 0.77 at 100 traversals, i.e. more non-place cells were classified as place cells as the number of traversals increased. The Combination method increased in sensitivity as the number of traversals increased from 2 to 20, above which it stabilised with a sensitivity of 0.73. The specificity of this method remained 1 regardless of the number of traversals included.

Thus our simulations predict that the Combination method would fail to detect at least 27% of hippocampal place cells, while up to 23% of the place cells identified by the Stability method would be false positives, and accuracy for both methods is affected by the number of times mice run through the environment.

### Place field properties

We next tested the effect of manipulating the width of the model place fields and their peak “fluorescence” on the ability of the three methods to detect the place cells (Fig 2C-F). First, we varied the width of the place field between 10 and 200 cm, equivalent to 5 to 100% of the total environment length, while keeping the peak value at a ΔF/F of 1.3. For both the Peak and the Stability method, varying place cell width did not materially affect sensitivity. However, the sensitivity of the Combination method was generally lower than the other methods, only approaching their sensitivity for place fields between 80 and 120 cm, and failing to detect any place field narrower than 20 cm or broader than 160 cm. Specificity of place cell detection was high for the Peak and Combination methods and unaffected by the place field width, whereas it was generally lower when using the Stability method (Fig 2D). It increased linearly with place field width for this method, presumably because the correlation between the fluorescence maps of the non-place cells and place cells decreases as the width increases.

**Fig 2.**
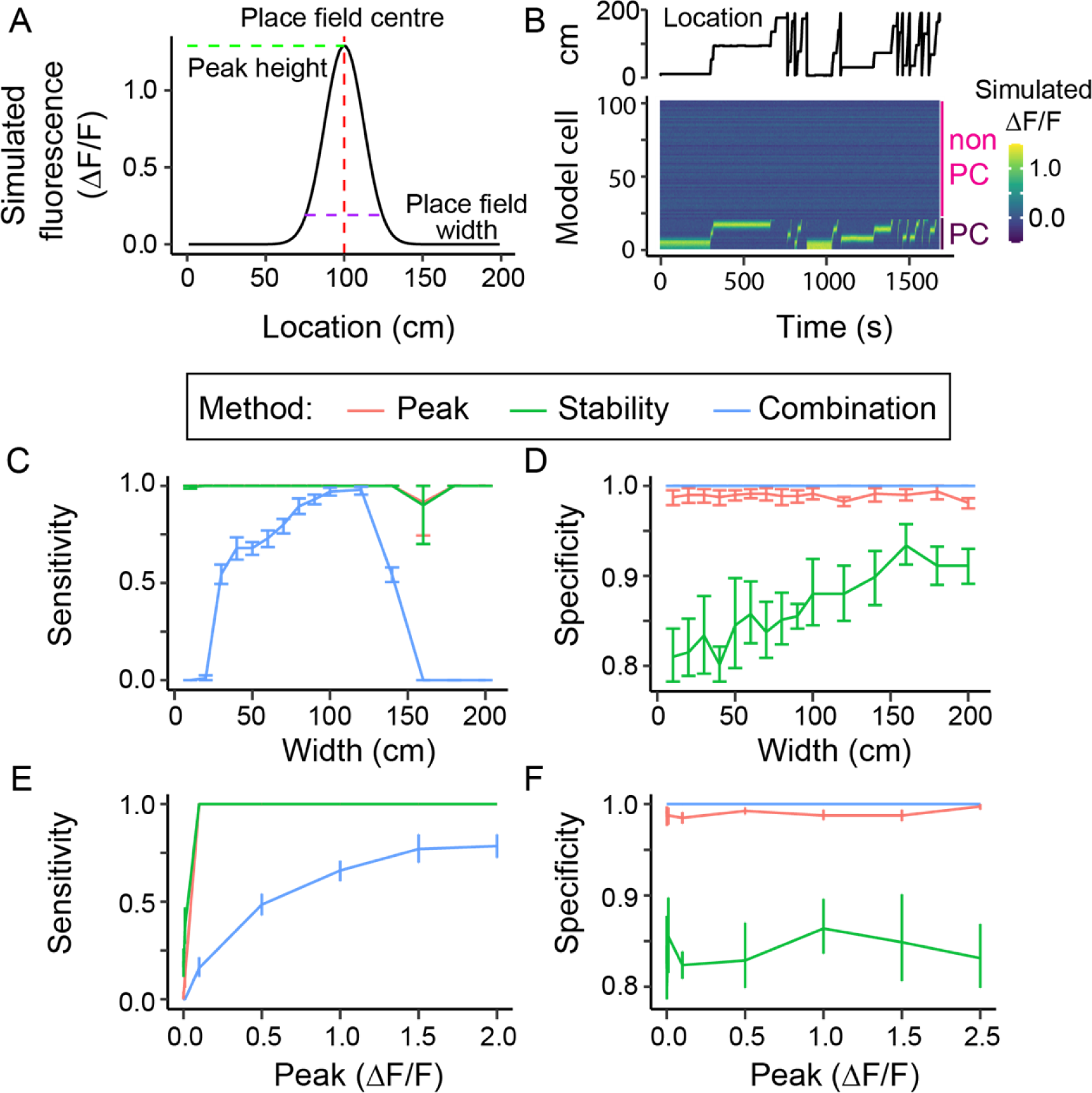
Place cell model and its effect on place cell detection. (A) Gaussian model of the fluorescence of a place field depending on the location (x-axis). The model takes the following parameters: the centre of the place field (red dashed line), the place field width (purple dashed line) and the peak of the Gaussian (green dashed line). (B) example fluorescence profile over time of a model data set containing 20 place cells (cells 1-20) and 80 non-place cells (cells 21-100), generated using the location trace shown. (C, E) Specificity and (D, F) sensitivity of the different methods as a function of the place field width (C, D) or peak (E, F). Data are means of 10 simulations. Error bars represent 95% confidence intervals.

We then varied the peak of the place field between a ΔF/F of 0.0001 and 2 (equivalent to 0.008% to 155% of the average fluorescence map peak measured in our two-photon data), keeping the place field width at 50 cm. Sensitivity was only affected using the Peak and Stability methods at extremely low, and therefore unrealistic, peak values (< 0.1, or <8% of peaks in our real data; Fig 2E). The sensitivity for the Combination method was generally lower and increased with place cell peak size. The specificity was not affected by changes in peak values for any of the methods (Fig 2F).

Next, we varied the number of place fields for each place cell, up to 4 place fields per cell (S3 Fig). Both the Peak and Stability methods were able to identify the model place cells, regardless of the number of place fields, with a high sensitivity, while the Combination method showed a dramatic drop in sensitivity above 2 place fields. Specificity was relatively unaffected by increasing the number of place fields, though was maximal using the Stability method when cells had 4 place fields.

Finally, we varied both the place field width and peak values at the same time (S4 Fig). The results reflect those when varying each parameter individually, illustrating the high general sensitivity of the Peak and Stability methods and the much narrower performance of the Combination method, which has high sensitivity at only a narrow range of widths and with high peak fluorescence values. Conversely specificity is high overall for the Peak and Combination methods but for the Stability method is lower across the range of parameters tested and lowest for narrow place fields.

In summary, the Combination method detected fewer model place cells and was the most affected by realistic variations in place field properties, while the Stability method classified more non-place cells as place cells and this tendency increased with narrower place fields. The Peak method was sensitive and specific over the whole range of place field properties tested.

### Variability and reliability decrease detection of model place cells, particularly by the Combination method

Because, physiologically, cells do not fire with identical profiles to repeated presentations of a stimulus [24], we simulated how the inherent variability of place cells affected their detection by the different methods. We manipulated the reliability of place cell responses by varying the percentage of traversals in which they fired (Fig 3A) and the location of the place field centre, by shifting the position of the activation peak with respect to the average firing field a percentage of the place field width (Fig 3B). The exact traversals that cells fired in were uniformly randomized, while the exact deviation of the place field centre was randomized using a Gaussian probability curve, and was repeated 10 times for each parameter.

**Fig 3.**
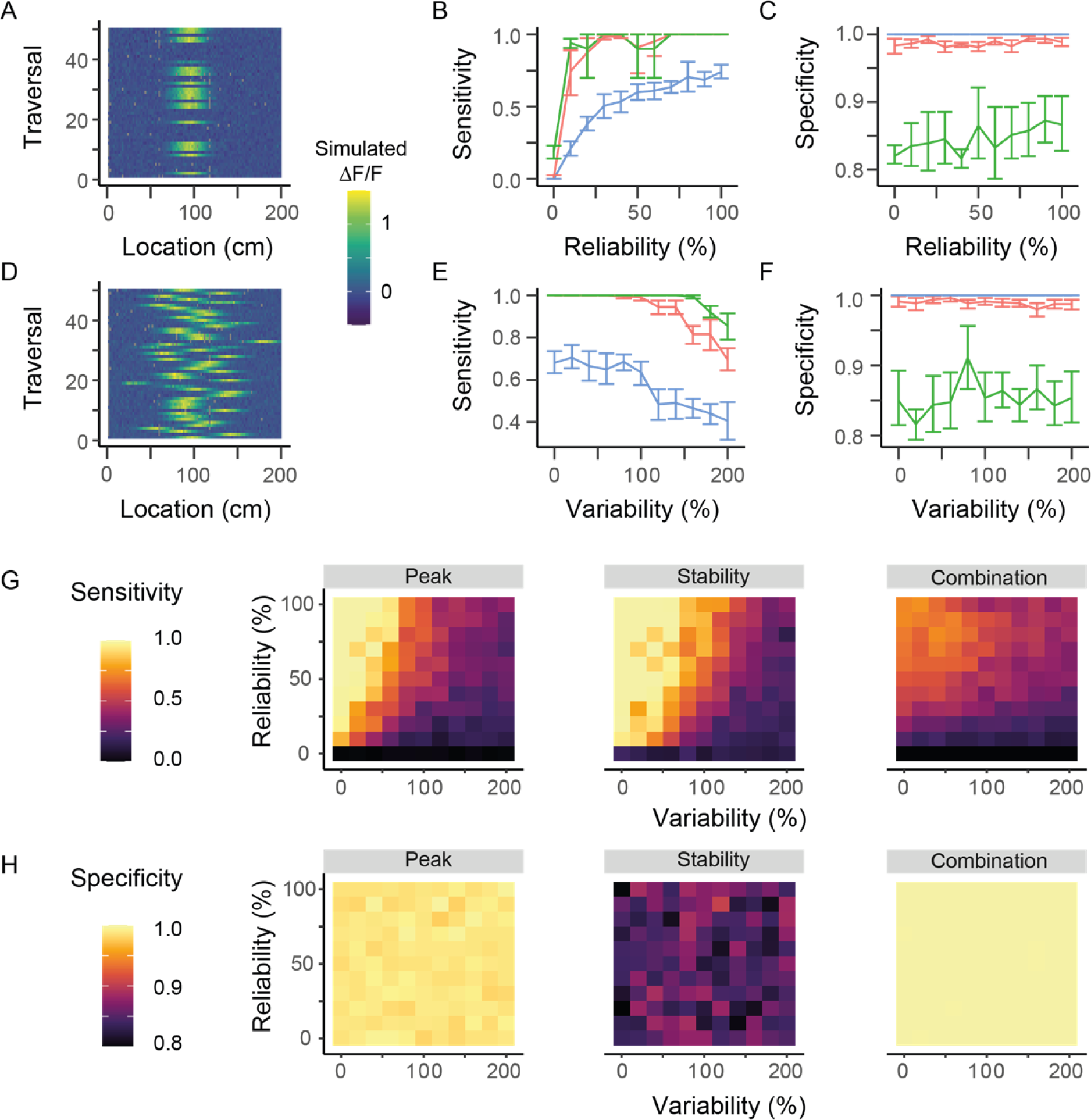
Place cell identification in variable and unreliable place cells. (A) Fluorescence map for a place cell with a reliability of 40% (B) Sensitivity of methods as a function of the reliability of a place cell. (C) Specificity of methods as a function of reliability. (D) Fluorescence map for a place cell with 60% variability. (E) Sensitivity of methods as a function of the variability of a cell’s place field across different traversals. (F) Specificity of methods as a function of variability. (G) Surface plots of the sensitivity of each method as a function of both reliability and variability. (H) Surface plots of the specificity of each method as a function of both reliability and variability. Grey points in A and D indicate the mouse did was not recorded at that location in that particular traversal, due to the limited frame rate of our recordings. Data shown are the means of 10 randomly created datasets, with the error bars representing 95% confidence intervals.

Both the Peak and the Stability method increased their sensitivity as cells became more reliable, reaching a maximum when cells fired in over 30% of traversals. The Combination method also increased in sensitivity with increased reliability, but at a slower rate, reaching a lower maximum sensitivity only when cells fired in 100% of traversals (Fig 3C). Varying reliability did not affect the specificity of any of the methods (Fig 3D).

Increased variability of the location of the place field centre caused a reduction of sensitivity above 100% variability for the Peak and Combination methods and above 150% variability for the Stability method (Fig 3E). Specificity of all methods was again unaffected by changes in variability (Fig 3F).

When changing both the variability and reliability, results were again similar to varying each individually, with maximal sensitivity for all methods occurring when cells had high reliability and low variability (Fig 3G), and stability being similar across all conditions (Fig 3H). Only the Peak method showed both high sensitivity and specificity.

In summary, the Peak method outperformed the other two methods when considering variable and unreliable responses. It could detect place cells when their firing properties were somewhat (but not very) unreliable or variable, suggesting it would cope with physiological variation in firing, and showed high specificity. Conversely, the Combination method’s sensitivity was decreased by even small decreases in reliability, while the Stability method detected even very variable cells, but had low overall specificity.

### The Peak method detects the most place cells in real datasets

The performance of the different methods of place cell detection on model data predicts that, on real datasets, the Peak method will have the highest sensitivity and specificity, with the Combination method being less sensitive and the Stability method often being less specific. This would suggest that in a real dataset, the Combination method would identify fewer place cells than the other two methods, while the Stability method, because it has a higher false positive rate on model data, might (inaccurately) detect more cells than the Peak method.

To test this, we collected neuronal calcium data from the stratum pyramidale of dorsal CA1 in 4 mice (8 imaging sessions in total) traversing through a 1D virtual reality corridor of 200 cm (Fig 4A). We applied each of the three methods to these data to identify which cells could be categorised as place cells (Fig 4B). There was a highly significant difference in the percentage of cells classed as place cells by the different methods (p =2.30x10^-4^, linear mixed-effects model), with the Peak method detecting the highest number (45.9 +/- 13.5% (mean +/- std)), followed by the Stability method (19.5 +/- 8.7%), while the Combination method only classified 0.9 +/- 1.0% of cells as place cells. The properties of these cells can be seen in an example recording (Fig 4C). Although no ground-truth exists for what should be included as a place cell, identified cells mostly seem to have clear place fields, suggesting classifications are appropriate, though in some cases cells are included that are somewhat non-classical (e.g. cell 13, Fig 4E, which has a spatial field in which it is not active).

**Fig 4.**
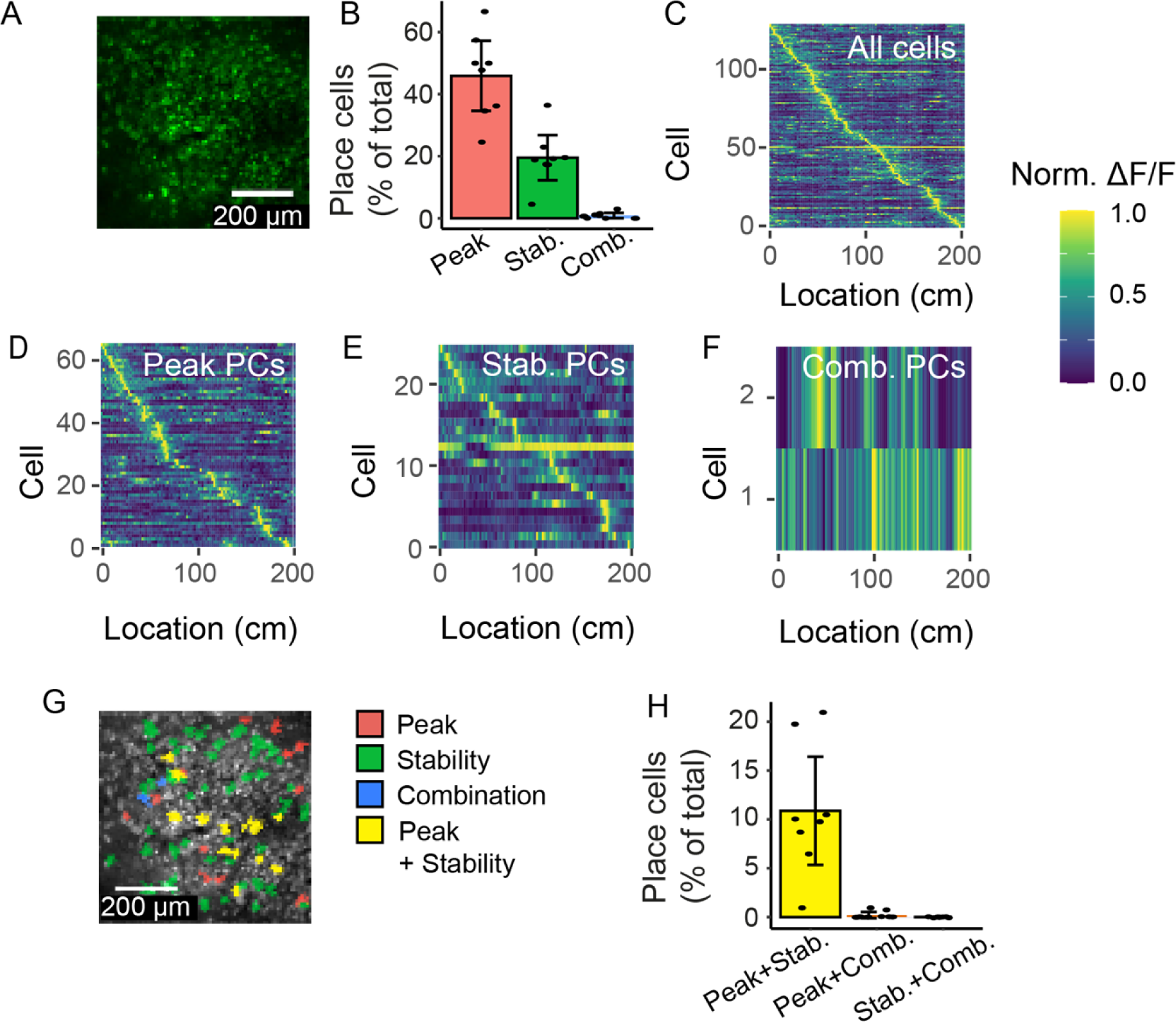
Place cell identification in real datasets. (A) Two-photon recording of GCaMP6f in pyramidal cells. Image is a standard deviation Z projection of a single recording. (B) Percentage of ROIs identified as place cells by the methods. (C) Fluorescence maps over location of ROIs in an example recording. Cells in this recording identified as place cells by (D) Peak, (E) Stability and (F) Combination methods. (G) Two-photon image from A showing ROIs identified as place cells by the three methods, including those identified by both Peak and Stability methods. (H) Comparison of percentage of ROIs identified as place cells by two methods. For B and H, each data point is a recording, bars show means, with error bars showing 95% confidence intervals.

### Differences between detection in modelled and real data

Based on the results from our modelled datasets, we expected the Stability method to classify more cells in a real dataset as place cells than the Peak method, but in fact it identified significantly fewer (p=2.45x10^-7^, Tukey post-hoc comparison). We probed the reasons for this difference both by modulating the properties of the modelled cells and inspecting examples of real cells identified differently by the different methods. Because the Stability method compares the activity of each cell to that of other cells in the population, while the Peak method compares each cell to a shifted version of itself, the properties of non-place cells could preferentially affect the performance of the Stability method. We therefore modulated the number and firing properties of the modelled non-place cells to see if this reduced the number of cells identified by the Stability method, but this could not account for the observed decrease in cells detected in real data (S5 Fig).

Visual inspection of the firing properties of cells identified by the Peak method, but not by the Stability method, however, showed that the difference in detection is caused, at least in part, by cells that either gain or lose a place field during the session, or that have different place fields at different times in the session (S6 Fig). The Stability method fails to identify these cells, because their mean activity over location for the first half of the session is different from the second. Minor changes to the Stability method or experimental design should prevent this effect, for example by comparing odd to even traversals, or by adding a familiarisation period for the animals at the start of each session which could ensure place fields are more stable when the experiment begins.

However, it is an open question whether some of the cells identified by the Peak method should really be classified as place cells, for example those with unstable place fields over the session.

### Different populations of real CA1 pyramidal cells are identified as place cells by the different methods

To understand whether the different methods simply identify more or fewer of the same population of cells as place cells, we determined the percentage of all ROIs that were identified by two methods. No cells were classed as place cells by all three methods. 10.9 +/- 6.6% of ROIs were identified as place cells by both the Peak and Stability methods (Fig 4H), while only 0.21 +/- 0.39% of ROIs were identified as place cells by the Peak and Combination methods and no cells were identified by both the Combination and Stability methods. These overlaps were not significantly different between pairs of methods, when corrected for expected overlap given the different percentages of cells identified by each method (linear mixed model on overlap vs method, with expected overlap as a covariate; p=0.53). Furthermore, the observed overlaps did not differ significantly from the expected overlap for the Peak-Stability and Peak-Combination comparisons (p=0.65, p=0.41, respectively, multiple Wilcoxon signed-rank test with Bonferroni-Holm correction). The overlap between Stability and Combination methods was below that expected by chance (measured overlap was 0%, expected was 0.19%, p=0.04). This suggests the methods are independent, and are not just identifying varying numbers of the same population of cells.

### Identified place cell populations differ on key characteristics

To better understand the nature of the different populations of cells identified by each method, we characterised their properties, comparing basic physical features, and functional characteristics. Place cells and non-place cells identified by the different methods were of similar sizes (Fig 5A; a smaller size could indicate a noisier image and a higher chance that the automated cell detection by Suite2P had made a mistake as identifying it as a cell). Place cells and non-place cells identified by the Peak and Stability methods were also similarly active, as assessed by both mean and maximum spiking rate (calculated from the fluorescence profile [25]), while place cells identified by the Combination method were less active than non-place cells (Fig 5B,C).

**Fig 5.**
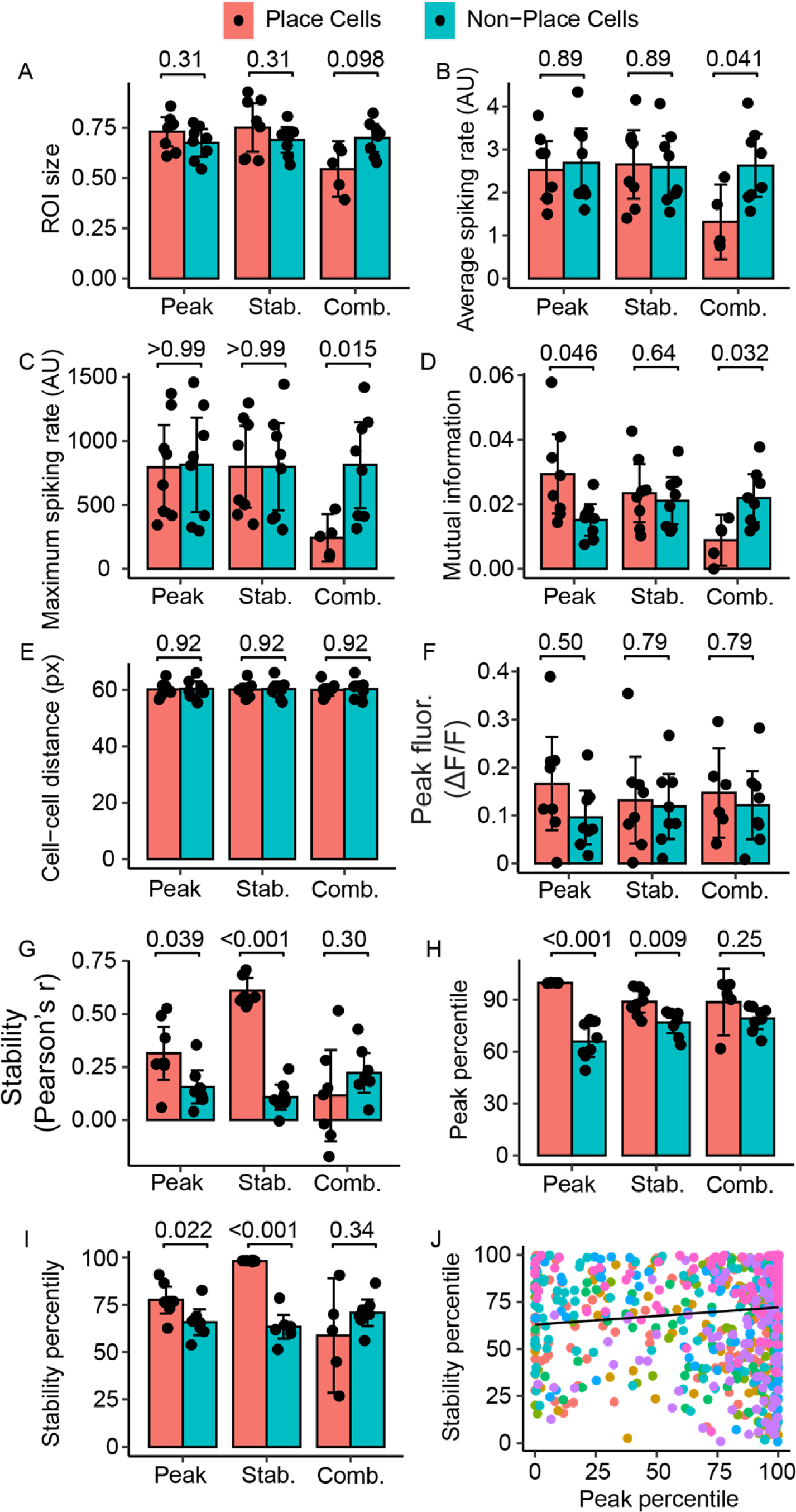
Characteristics of place cell populations. (A) The ROI size, as a percentage of the size of the largest ROI in that recording, of the place cell and non-place cell population in the three methods. (B) Mean spiking rate of place cells and non-place cells. (C) Maximum spiking rate of place cells and non-place cells. (D) Mutual information between cell activity and location for each classification by the three methods. (E) Mean distance between the centroids of the place cells (pink), and the distance from place cells to non-place cells (blue). (F) Peak intensity in the fluorescence maps, the main characteristic used in the Peak method. (G) Intrasession stability, measured using the Pearson correlation coefficient, between the fluorescence map of the 1^st^ half and the fluorescence map of the 2^nd^ half for place cell and non-place cell, the main characteristic used in the Stability method. Percentile of (H) peak intensity and (I) intrasession stability compared to shuffled controls. (J) Correlation between percentile of peak intensity and intrasession stability for all cells. In J, each colour indicates a single dataset. For the other figures, bars are the means of 8 datasets, black dots are values for each individual dataset and error bars represent the 95% confidence interval. P-values for A-I are from multiple paired t-tests with Bonferroni-Holm correction.

To investigate the information about location carried in place or non-place cells identified by the different methods, we calculated the mutual information between the cell’s activity and the animal’s location, providing an approximation of the spatial information carried in each cell’s calcium signals [26–28]. More spatial information was carried in place cells identified by the Peak method compared to non-place cells, but there was no difference in mutual information held by the place cell and non-place cell populations for the Stability method (Fig 5D). Furthermore, place cells identified by the Combination method carried significantly lower spatial information than non-place cells, presumably because cells that the other methods would call place cells are classified as non-place cells using the Combination method. In addition, cells may convey information about location without being a place cell. For example, a cell whose activity ramps up gradually as the mouse progresses through the environment does not have a clear place field and will not be included as such in these methods but does communicate substantial spatial information.

Place cells exhibit no particular spatial organisation [10], and any such organisation compared to non-place cells might indicate an unwanted selection by one of the methods, e.g. based on illumination levels in one part of the field of view. Therefore, we determined the physical location of the detected place cells within the field of view of the recording. We expect the place cells and non-place cells to be randomly distributed within the field of view, as previously reported [10]. We determined the location of the centroid of each cell, labelled them as either place cell or non-place cell according to the different methods and then determined the distance from each place cell to all other place cells, and each place cell to all other non-place cells (Fig 5E). Place cells were no more similar in location to other place cells than to non-place cells for any of the methods, suggesting that place cell detection is not being influenced by position within the field of view.

### Dependence of place cell detection on model parameters

The lack of overlap between cells identified by the different methods suggests that they identify fundamentally different populations of cells as being place cells, i.e. there is not a single population of cells that fits all the different definitions of place cells used by the different methods. To explicitly test for this, we calculated the extent to which the parameters which underlie classification for the Peak and Stability (peak intensity and correlation in firing across the session, respectively) were represented in the groups of place cells that were selected by each other method (The Combination method uses several parameters so could not be analysed in this way). None of the methods had significantly higher peak intensity in the place cell group compared to the non-place cells (Fig 5F), including the Peak method.

Place cells identified by both the Peak and the Stability methods, but not the Combination method, had an increased correlation in activity across the session in place cells compared to non-place cells – i.e. they were more stable (Fig 5G). Thus, the Peak method also selects for stable cells – likely because stable place cells are consistently active in the same location across traversals, yielding on average a robust peak.

Though both the peak intensity and stability are parameters used as a measure to identify place cells in their respective methods, they compare the values of each individual cell to the peaks of shuffled versions of that same cell, rather than comparing it to the wider population. This means that place cells are not necessarily the cells with the highest absolute peaks or highest correlation, but rather the highest value relative to their own shuffled controls. Therefore, we calculated the percentile these values fell in, relative to the shuffled controls, for each cell. Place cells identified by both the Peak and the Stability methods, but not the Combination method, had an increased percentile score for both the stability and the peak parameters (Fig 5H,I).

To understand why the Peak and Stability methods select place cell populations with similar peak intensity and stability, we determined to what extent the parameters used to define place cells were independent from each other. We correlated the percentiles for the peak intensity and intra-session correlation for each cell across all datasets (Fig 5J). There was a small but significant correlation between the percentiles for peak fluorescence and intra-session correlation (Pearson correlation coefficient, R=0.08, p =0.013). Although this suggests the two measures are not completely independent – cells with high peak values have a tendency to have more highly-correlated firing patterns within a session – we did not see this reflected in the overlap between the methods (Fig4H, S7H Fig). This may be because the correlation coefficient between peak intensity and intra-session correlation is too low to have an impact. In addition, it may be the case that more correlated cells are not that same as those that reach the threshold for inclusion as a place cell in one or both methods, so are not reflected in the place cell overlap.

### Optimisation of the Combination method

The Combination method identified many fewer cells in our data than in previously publications [10, 19] or the other two methods applied here. We wondered if this could be due to the number of thresholds the Combination method applies, ([10, 19] code to implement the Combination method shared by the authors) which implement the Combination method, requiring putative place cells to fall within specific parameter ranges. These were presumably optimised for the published datasets and may make the method less transferable to other datasets. To test this, we investigated how varying these parameters and thresholds affected place cell detection in our model and experimental datasets (S7 Fig; Methods). Sensitivity for detecting model place cells could be improved by increasing the threshold for place field fluorescence compared to baseline (S7B Fig) or reducing the threshold in-field/out-field fluorescence ratio to 2 (S7E Fig), but neither modification caused a substantial increase in the number of cells classed as place cells in experimental data (1.2%; S7G Fig). Instead, the critical feature of the Combination method is the bootstrapping method, which splits the fluorescence data into chunks and shuffles them. Cells are only classed as place cells if fewer than 5% of that cell’s shuffles also fit the classification criteria. If we replaced the bootstrapping methods from the Combination method with that of the Peak method, which instead offsets fluorescence traces relative to time, causing more disruption to the association between fluorescence and location, we identified 15% of ROIs in our experimental data as place cells (S7G Fig), which is within the published range using the Combination method [10], though below the numbers identified by the Peak method in our data (Fig 4B). Some of these newly identified cells now overlapped with those identified by other methods, 11.5% of all ROIs now overlapping with cells identified by the Peak method and 3.9% overlapping with those identified with the Stability method (S7H Fig). 3.3% of ROIs were identified as place cells by all methods. Therefore, when using the new bootstrapping method, around 75% of PCs identified by the Combination method were also identified by the Peak method. However, these levels of overlap were still as expected given the frequency of detection by each method (p=0.92 for all comparisons: Peak-Stability, Peak-Combination and Stability-Combination;, multiple paired t-tests of expected vs. observed overlap, with Bonferroni-Holm correction). This means that the cells identified by each method are still likely to be from independent populations (i.e. if a cell is classified as a place cell by one method that does not increase its chance of being classified as a place cell by another method).

## Discussion

We compared three different methods for detecting place cells in two-photon calcium imaging data from mice running through a one-dimensional environment, comparing sensitivity and specificity of the different methods to model cell data, and whether they identified the same real populations of cells as place cells. The methods performed very differently at detecting model place cells and were differentially sensitive to changing parameters (e.g. number of traversals of the environment, place cell width, consistency of firing). This suggested that variability in firing properties across real hippocampal CA1 pyramidal cells would lead to different cells being classified as place cells by the different methods. This proved to be the case: there was very little overlap in real CA1 place cell populations classified by the different methods, because the key features used by the different methods to classify place cells proved only weakly correlated. When research groups use these different methods to detect place cells, they therefore identify and study largely independent populations of cells. Researchers should therefore explicitly consider the properties they are selecting for when choosing a place cell detection method, as this decision will determine which cells they do and do not select. In the future, use of a single method across different groups would better allow comparison of results across the field. The increased selectivity and specificity of the Peak method when using model data and the increased spatial information carried in place cells compared to non-place cells, leads us to conclude that the Peak method will usually be the best for detecting place cells in calcium imaging data. Nevertheless, some curation of detected cells may still be desirable, as some cells identified by the Peak method showed unstable firing patterns that may be considered too dissimilar to the firing properties of a traditional place cell.

### What is a place cell?

Because our results reveal that the choice of detection method will determine the population of cells identified as place cells by a given experiment, it is important to consider what the “desirable” properties of a place cell are, and whether the different classification methods detect these properties. O’Keefe’s first description of place cells defined them as “those for which the rat’s position on the maze was a necessary condition for maximal unit firing.” [1]. Of the three methods we assess here, the Peak method comes perhaps closest to this original definition, requiring cells to have high and consistent firing in one location compared to others, by statistically comparing whether the peak response consistently occurs at the same location. The Combination method achieves something similar by requiring firing to be above a certain threshold for a defined range of place field sizes, while the Stability method only requires the firing pattern to be stable over location and time without the explicit requirement for the peak to be highest in one location.

The reason for the divergence of place cell definitions from the initial more constrained description is the increased emergence of variability in “place cell” properties. We now know that the size, shape, number and stability of place fields are not constant but can be affected by the environment and where in CA1 they are recorded: Place fields are wider, and more numerous, in larger environments, more stable in the presence of cues and smaller and less spatially selective in dorsal vs. medial CA1 [29–33]. Clearly methods that discriminate place from non-place cells according to a rigid interpretation of the “classic” definition will miss cells with these more variable properties. Indeed, we found that the Combination method has a lower sensitivity to detect cells with a place field outside a narrow range (100-120 cm), and failed to detect place cells as variability of place field position increased or reliability of firing reduced. Thus, we predict that experimental manipulations that alter the size of the environment and number of cues provided would reduce the number of place cells identified using the Combination method, but not using the Peak method.

Similar changes in place cell properties can occur due to the method of acquiring neural data, impacting on which might be the most appropriate method to choose to detect these cells. Many experiments that record the activity of large populations of cells, either using calcium imaging, as here, or multi-channel electrode arrays are conducted in head-restrained mice, while the animals are running on a 1D linear track [10,34–36]. These conditions likely cause place cells in our experiments to have broader place fields than in real world environments [14, 37], and to be more directionally-sensitive, due to the 1D nature of our linear track [10, 38]. The place cell detection method used for such imaging data therefore needs to recognise these less sharply tuned activity patterns. We found that the Peak method detected the most place cells in our experimental data, and this was the only method in which the detected place cells carried a higher mutual information for location than non-place cells. This suggests that only the Peak method was able to classify enough of the spatially-selective cells in our head-restrained imaging data as place cells, perhaps because of its broader spatial selectivity compared to classic tetrode recordings, and suggests that similar studies using head-fixed mice and linear tracks would also miss spatially-selective cells if a more conservative method were employed.

Less controllable variations in experiments between labs, for example the amount the mice run in an experiment, will also differentially affect the number of cells classed as place cells by the different methods. Our simulations suggested that all three methods would be more sensitive at detecting place cells as mice made more traversals of the environment, but the Combination method would miss cells with even relatively high numbers of traversals (missing over 50% of cells when 10 traversals were made). Furthermore, the Stability method increasingly falsely characterised non-place cells as place cells as the number of traversals increased. Thus, only the Peak method could correctly identify place cells with low or high numbers of traversals. Choice of place cell categorisation method will therefore clearly affect the number of cells identified in experiments where animals run to different degrees. Crucially, however, the confounding of place cell identification with running behaviour in 2 of the 3 methods could lead to inaccurate conclusions about how experimental manipulations affect place coding. If, for example, mice of a certain genotype run less, generating fewer traversals, they could appear to have fewer place cells using the Combination method, or more place cells if using the Stability method, without the number of “true” place cells having changed.

In this study, we only investigated the effect of varying the number of traversals on place cell detection, but other characteristics of the locomotion may also have an effect, for example running speed or the amount of starting and stopping. As these effects may interact with the environment or task, (for example, mice slowing down before a reward delivery), we suggest testing the performance of a chosen method of place cell detection on a model dataset with the expected locomotion characteristics before using it to detect place cells experimentally. We provide source code (https://github.com/DoriMG/place_cell_methods) to facilitate this process, into which experiment-specific patterns of locomotion can be loaded and used to predict place cell detection using the three methods described here.

In conclusion, because place cells have a more variable activity pattern than was originally thought, particularly across the large populations and range of experimental paradigms permitted by calcium imaging, classification methods should be sufficiently able to identify cells that vary in terms of the size, and reliability and variability of their place field location. However, the different methods we tested select largely different populations of cells which differ in key characteristics, highlighting that choice of place cell classification method is critical for the conclusions a study will draw as to the nature of place cells. We provide model place cell code to help researchers test their how their chosen methods or experimental manipulations might affect detection and the properties of place cells in their data. However, we suggest that consensus in the field for an identification method would help inter-study comparability. Overall, we found that the Peak method, which has previously only been used on electrophysiology data, showed the best results on two-photon calcium imaging data out of the three tested methods. It demonstrated a high selectivity and specificity for selecting model place cells that was robust to moderate changes in place field properties but decreased appropriately as the reliability and variability for the place field decreased. It detected more place cells in a real dataset and these cells carried more mutual information about location than non-place cells, unlike the other methods. For most experimental designs, we therefore recommend use of the Peak method for classifying place cells in calcium imaging data.

## Materials and Methods

### Ethics Statement

Experiments were approved by the UK Home Office, in accordance with the 1986 Animal (Scientific Procedures) Act as well as the University of Sussex Animal Welfare Ethical Review Board.

### Animals

Experiments used four C57/BL6 mice (2 female, 2 male) expressing the genetically-encoded calcium indicator GCaMP6f under the control of a Thy-1 promoter (C57BL/6J-Tg(Thy1-GCaMP6f)GP5.5Dkim/J). The mice were housed in a 12h reverse dark/light cycle environment at a temperature of 22°C and were given ad libitum access to food and water.

### Hippocampal cranial window surgery

Surgery was performed when mice were a minimum age of eight weeks. Before surgery, mice received subcutaneous injections of dexamethasone (60 μL, 2mg/mL), saline (400 μL) and buprenorphine (40 μL, 0.3mg/mL diluted 1:10 in saline) to reduce inflammation, for hydration, and pain relief respectively. Mice were maintained at 0.8-2.0% isofluorane anaesthesia for the duration of the surgery. Body temperature was maintained at 37 °C using a homeothermic blanket (PhysioSuite, Kent Scientific Corporation). A craniotomy was inserted above dorsal CA1 as previously described [23]. Briefly, the skin above the skull was removed and the skull was scored to increase the surface area for binding dental cement. A custom-made stainless steel headplate was then fixed to the skull with black dental cement (Unifast Powder mixed with black ink (1:15 w/w) and Unifast Liquid). A 3 mm diameter craniotomy was then performed 2mm posterior to bregma and 1.5 mm lateral to the sagittal suture. Following the removal of the skull flap and the dura, brain tissue overlying the hippocampus was aspirated (New Askir 30, CA-MI Srl) until vertical striations of the corpus callosum were visible. We then inserted a custom 3D printed cannula (2.4 mm ID, 3 mm OD, 1.5 mm height) made of a biocompatible Dental SG resin (FormLabs) so that the glass coverslip at the bottom of the cannula lay directly on top of the brain tissue. The top of the cannula had a rim (0.2mm height, 3 mm OD) resting on top of the skull, which was attached using tissue adhesive (3M VetBond) and then covered with more dental cement. A rubber ring was then attached on top of the headplate for subsequent use as a well for the water needed for the water-immersion microscope objective. The mice were given an injection of meloxicam (125 μL, 5 mg/ml) as an analgesic near the end of the surgery and then received meloxicam (200 μL, 1.5 mg/mL) for 4 days following the surgery via oral admission. Their health was monitored and they were weighed daily.

### Two-photon imaging

#### Habituation

A week or more after surgery, the mouse was habituated to the imaging rig by head-fixing it for an increasing amount of time each day for at least a week before it was imaged. During the habituation it was also presented with the virtual reality environment several times to make it familiar with this setup.

#### Imaging rig (S2 Fig)

The mice were head-fixed above a polystyrene cylinder, on which they could run. The cylinder was fitted with a rotary encoder (Kübler, 4096 pulses per revolution). Two screens in front of the mice were used to display a custom virtual reality (VR) environment designed using ViRMEn [39].

#### Stimulus presentation

The virtual reality environment presented to the mice was a wide corridor, 200 cm long and 80 cm wide, with patterned walls (30 cm high) and floor (see S2 Fig). Three sets of objects (spatial cues) were present outside the walls: blue square pillars with white stripes, light blue cones with white and gray diagonal stripes and grey cylinders with green dots. The objects were placed at 65 cm, 140 cm and 200 cm from the start of the corridor. All objects were 100 cm high and visible from the start of the arena. Both before and after the wide corridor was a dark grey tunnel with a diameter of 30 cm, 50 cm long before the corridor and 45 cm long after the corridor) which served to allow smooth transitions between multiple presentations of the environment. The mice were not required to perform a task while in the virtual environment.

#### Data acquisition

The stratum pyramidale of dorsal CA1 was imaged using a two-photon microscope (Scientifica) with a water-immersion objective (CFl75 LWD 16X W, Nikon; 0.80 numerical aperture, 3 mm working distance). GCaMP6f was excited using a Chameleon Vision II Ti:Sapphire laser (Coherent) at a wavelength of 940nm with a gallium arsenide phosphide photomultiplier tube. We used the ScanImage software (Vidrio Technologies, MATLAB) to control the microscope and collect data. The stratum pyramidale was identified from the presence of densely packed cell bodies. Image acquisition used a wide field-of-view (547 x 547 μm) at a low resolution to optimise the acquisition rate (128x128 pixels, 7.51Hz, pixel size 4.27 μm). Sessions lasted between 44 and 45 minutes.

### Image Analysis

#### Preprocessing

Preprocessing was conducted using Suite2P software [25]. Firstly, images were registered using the default settings, then regions of interest (ROIs) corresponding to pyramidal cell bodies were identified based on their morphology (having a diameter of approximately 2 pixels/8.5 μm) and a tau, the decay time for the calcium indicator, of 0.8. We trained a classifier by manually curating the detected ROIs based on the mean image of the original recording, the shape and the activity pattern of the ROI. On average 58 +/- 4% of ROIs were excluded per imaging session. We obtained the calcium signal corrected for neuropil activity for each ROI from the Suite2P output.

For each ROI, fluorescence time courses were normalised to baseline fluorescence by dividing the whole trace by the average intensity in that ROI during the first 100 frames of the recording. For calculations of fluorescence maps, any frames where the mouse was stationary were excluded (defined as the speed being below 10 % of the maximum speed).

An extra pre-processing step was used in Dombeck et al. [10] and associated papers, described in full in [11], so we at first used this step as well when replicating this method (hereafter referred to as the Combination method). After ROI extraction and normalisation the whole time series was divided by the baseline, which was defined as the 8^th^ percentile of values in each ∼15 second interval. Significant transients were then identified as calcium events that started when fluorescence deviated more than 2 standard deviations from the baseline and ended when the fluorescence returned to less than 0.5 standard deviations from the baseline. Fluorescence outside of the significant events was set to 0. However, we noticed that this preprocessing step led to place cell activity appearing negative, and thus these cells being rejected, if their activity was shorter than 15s and occurred in the presence of a negative baseline. For our data, this led to all cells being rejected at this stage. We therefore did not use this preprocessing method in our analyses.

Subsequently, to test whether the lack of preprocessing explained the low number of cells identified using this method, we amended the preprocessing method to correct for slow drifts in the baseline while preventing division by a negative baseline, by subtracting rather than dividing by the baseline.

This caused a small but significant increase in the sensitivity of the Combination method compared to non-preprocessed traces (S7A Fig).

### Model data generation

We generated model place cells to test the performance of the place cell detection methods. Time series of calcium responses of model cells were generated using real locomotion traces of mice running through a linear virtual environment. 8 datasets containing 184 traversals were split into separate traversals and model locomotion traces were generated by randomly selecting the required number of traversals (each traversal could be selected more than once). Model place cell properties (described below) were convolved with these locomotion traces to generate a fluorescence map for each cell, with the same “frame rate” as used in the real imaging sessions used to acquire locomotion (7.51 Hz).

For each dataset, we included 20 model place cells – in which fluorescence was modulated by spatial location – and 80 non-place cells – whose fluorescence was independent of location. This ratio was chosen to mimic the percentage of place cells typically reported in experiments using a one-dimensional track and two-photon microscopy [10]. The inclusion of non-place cells was crucial as some of the place cell detection methods use comparisons with other cells in their definition, and thus rely on the population containing non-place cells as well as place cells.

For the model place cells each place field was modelled as a 1D Gaussian field centred at a randomly selected location (Fig 2A). The sigma of the Gaussian was set to 12.5 cm, such that 95% of values – 4 times sigma – fall within the place field width of 50 cm, the average place field width reported by Dombeck et al. [10]. The peak of the Gaussian (1.3) was determined using the top 10% of each cell’s fluorescence in its fluorescence map from 992 cells from 8 datasets.

The noise in all cells was modelled using a Poisson distribution (generated using the MATLAB poissrnd function), with lambda estimated from the raw traces of 992 cells from 8 datasets. The median lambda of the 10% of cells that had the best Poisson fit (235.1, as estimated from the sum of squared errors; SSE) was used as the lambda for our model noise. Raw noise was generated using this parameter, then ΔF/F of this noise was calculated to obtain the normalised noise used in the model cells. The average ΔF/F of the noise, as tested on 10000 traces each with a length of 1000, was 0.0024 +- 0.0467, which is 0.18% of the peak value.

Non-place cell traces usually consisted of only noise (Fig 2B, top 80 cells), generated as above, while the noise was added to the generated place field for the place cells (Fig 2B, bottom 20 cells). For S5 Fig, we also considered the impact of non-spatial firing of non-place cells. In our real data, there were on average 0.0062 peaks per frame, i.e. a calcium peaks occurred about every 160 frames. We therefore randomly added a range of peaks to non-place cell traces, from 0- 0.02 peaks per frame.

### Manipulation of place cells

We manipulated the place cell properties in order to model the imperfect nature of real place cells. We varied the place field width, reliability, spatial variability and the number of place fields per place cell.

#### Place field width

The place field width was varied by varying the sigma of the Gaussian field for each place cell.

#### Place field peak

The place field peak was varied by scaling the Gaussian model using a single scalar. This way, the shape of the Gaussian remained, while the overall peak could be increased or decreased.

#### Number of place fields

We varied the number of place fields between 1 and 4, spaced to cover the whole environment but with no overlap. The upper limit of 4 place fields was therefore set by the size of the place field (50 cm) and the environment length (2 m).

#### Reliability

We defined the reliability of a place cell as the probability it will have a place field in a given traversal. The probability P_field_ was between 0 and 1, where P_field_ = 0 meant the place cells did not have a place field in any traversal, and P_field_ = 1 meant the place cells had place fields in every traversal. At a probability between the 0 and 1 the place cells had place fields in a randomly selected proportion of traversals equal to P_field_ times the total number of traversals.

#### Spatial variability

We defined the variability of a place cell as the average deviation of the place field per traversal from the centre of the average place field. We modelled this by defining a Gaussian centered around the centre of the average place field, with a flatter Gaussian (i.e. higher sigma) equating to a higher variability. For each traversal we drew a value from this Gaussian distribution to be the centre of the place field for that traversal.

### Performance measures

To calculate the performance of the place cell detection methods, we calculated the sensitivity and specificity of each method using the number of true positives (TP), true negatives (TN), false positives (FP) and false negatives (FN). The sensitivity is a measure of how well the method is able to find all the true place cells in the dataset, while the specificity is a measure of how specific the method is for identifying just true place cells without false positives.

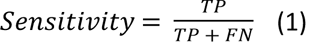

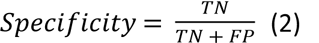

### Place cell detection methods

We tested three methods for place cell detection (S1 Fig) all of which have all been used in previous studies to identify place cells.

#### Peak method

This method of place cell detection was described for electrophysiology data from mice running through a 1D virtual corridor [20]. We adapted the original method for use with fluorescence data as follows: We first calculated the fluorescence maps (the average fluorescence in each location bin) for each cell from which we determined the peak fluorescence for each cell. The neuronal fluorescence data was then randomly shuffled 500 times relative to the location data by shifting the fluorescence data in time randomly by at least 5 seconds. Because the running speed of the mouse is not constant, this alters the fluorescence at each location and therefore the “peakiness” of the fluorescence map. Each cell’s fluorescence map and peak fluorescence was then determined for each shuffle. Any cell with a true peak fluorescence in the top 1% of all shuffles was deemed a place cell. Thus, the peak method detects cells with high fluorescence in a place field compared to the baseline.

#### Stability method

The stability method was developed by O’Leary [21] and is based on classic methods of detecting place cells in real-world two-dimensional (2D) environments which require place cells to have a consistent place field over multiple traversals (e.g. [3]). First, a separate fluorescence map was calculated for each cell for each of the two halves of a recording session and the linear correlation between these two fluorescence maps was calculated. These correlation coefficients were then compared to control values obtained by comparing the fluorescence map of the first half of a session for that cell with the fluorescence maps of the second half of other randomly selected cells in the dataset (100 repeats per cell with redraws). The within-cell correlation of the cell was compared to all the shuffles for that cells, and if the correlation was above the 95^th^ - percentile of the shuffles, it was deemed a place cell.

#### Combination method

The Combination method [10,18,19] uses a combination of the level of fluorescence above baseline, with stability of firing over different traversals. It uses pre-processed fluorescence traces of cells that have been thresholded so they only include significant transients, followed by a number of thresholding steps to find “true” place fields. This method has been modified over the three publications cited above, the most recent of which was used here. First, the fluorescence map was calculated from the preprocessed traces for each cell and possible place fields were determined by thresholding the fluorescence map with a cut-off value of 0.25 times the difference between the peak fluorescence value and the baseline. Possible place fields were defined as having above-threshold fluorescence in contiguous locations for at least 20 cm and less than 120 cm. Each field was also required to have one bin with a value of at least 10% of the mean fluorescence of that cell over the session. The mean fluorescence in each place field was then divided by the mean fluorescence outside of the place field, and cells were only deemed place cells if the in-field/out-field ratio was higher than or equal to 4. Lastly, the cells were required to have a significant transient, as defined in the pre-processing thresholding step, in at least 20% of the traversals of the environment. All traces were shuffled 1000 times with respect to location and classified using the above methods. Only cells that were classified as place cells in fewer than 5% of the shuffles were considered to be true place cells. Thus, the Combination method requires place cells to significantly increase fluorescence from baseline over a contiguous place field, with some stability of responding over time.

### Statistical analysis

All statistical tests were conducted in R [40]. Where appropriate, a Shapiro–Wilk test was used to test the data for normality. If the data was determined not to deviate from a normal distribution (p>0.05), we performed the appropriate parametric test, otherwise a non-parametric test was applied. The tests performed for each comparison are detailed in the text.

### Data and software availability

The MATLAB used to automatically generate model datasets is available on GitHub (https://github.com/DoriMG/place_cell_methods). This repository also contains the locomotion dataset we used to generate the model data. We further provide an explanation of how to use the code to generate novel model datasets with varying parameters. Experimental data used for each relevant figure is available via FigShare (10.6084/m9.figshare.13560548).

## Acknowledgements

With thanks to Alice O’Leary and Daniel Dombeck for sharing their code. We would like to thank O. Hall-Bird and M. Hall-Bird for their insights and useful suggestions on this project.

---

**S1 Fig.**
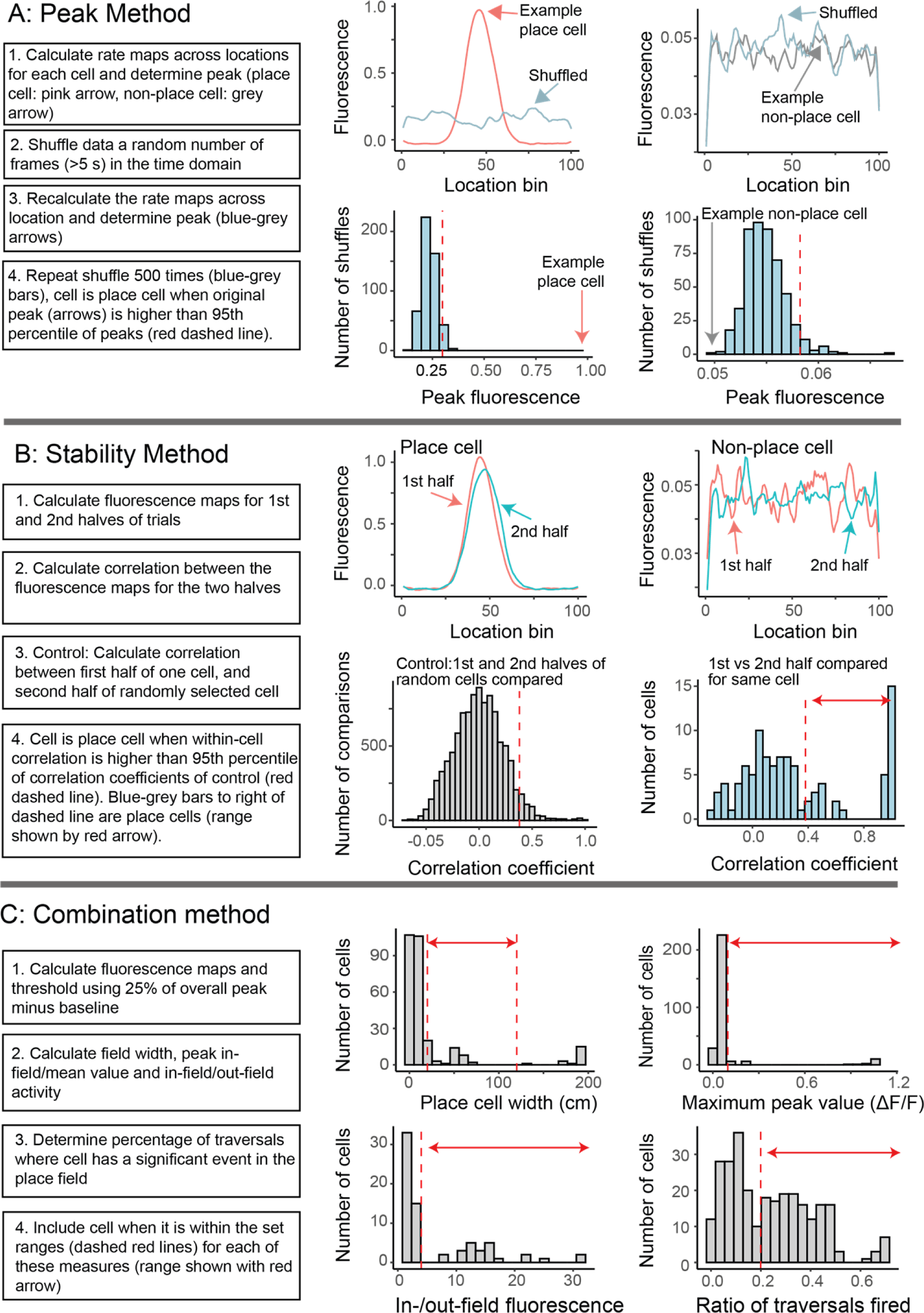
Graphical summary of tested methods. A graphical representation of how the (A) Peak, (B) Stability and (C) Combination method define a place cell based on its fluorescence map.

**S2 Fig.**
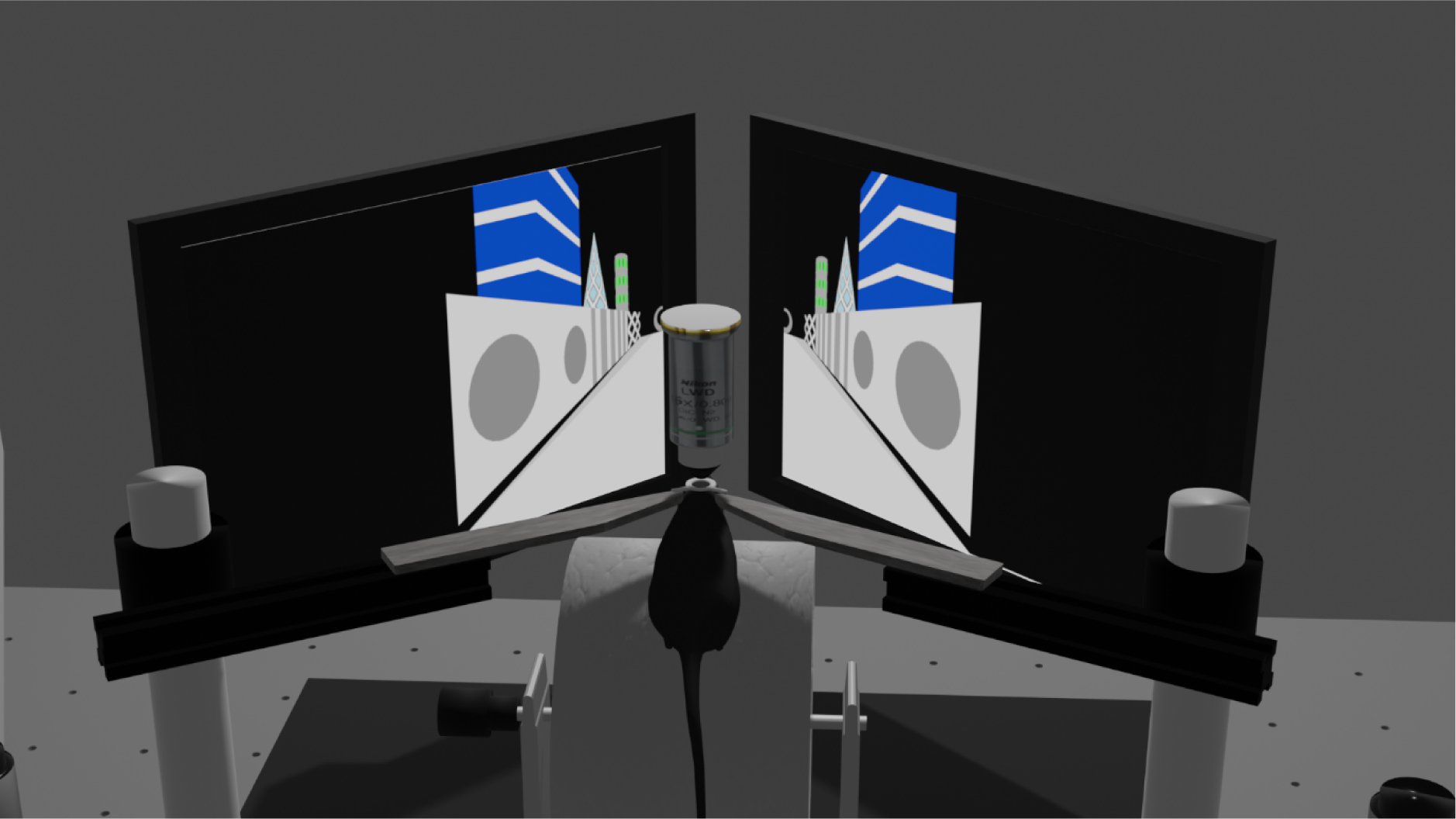
Experimental setup. The two-photon experimental setup as seen from behind the mouse. The mouse is head-fixed and standing on a wheel that it can use to control the environment projected onto the screens in front of it.

**S3 Fig.**
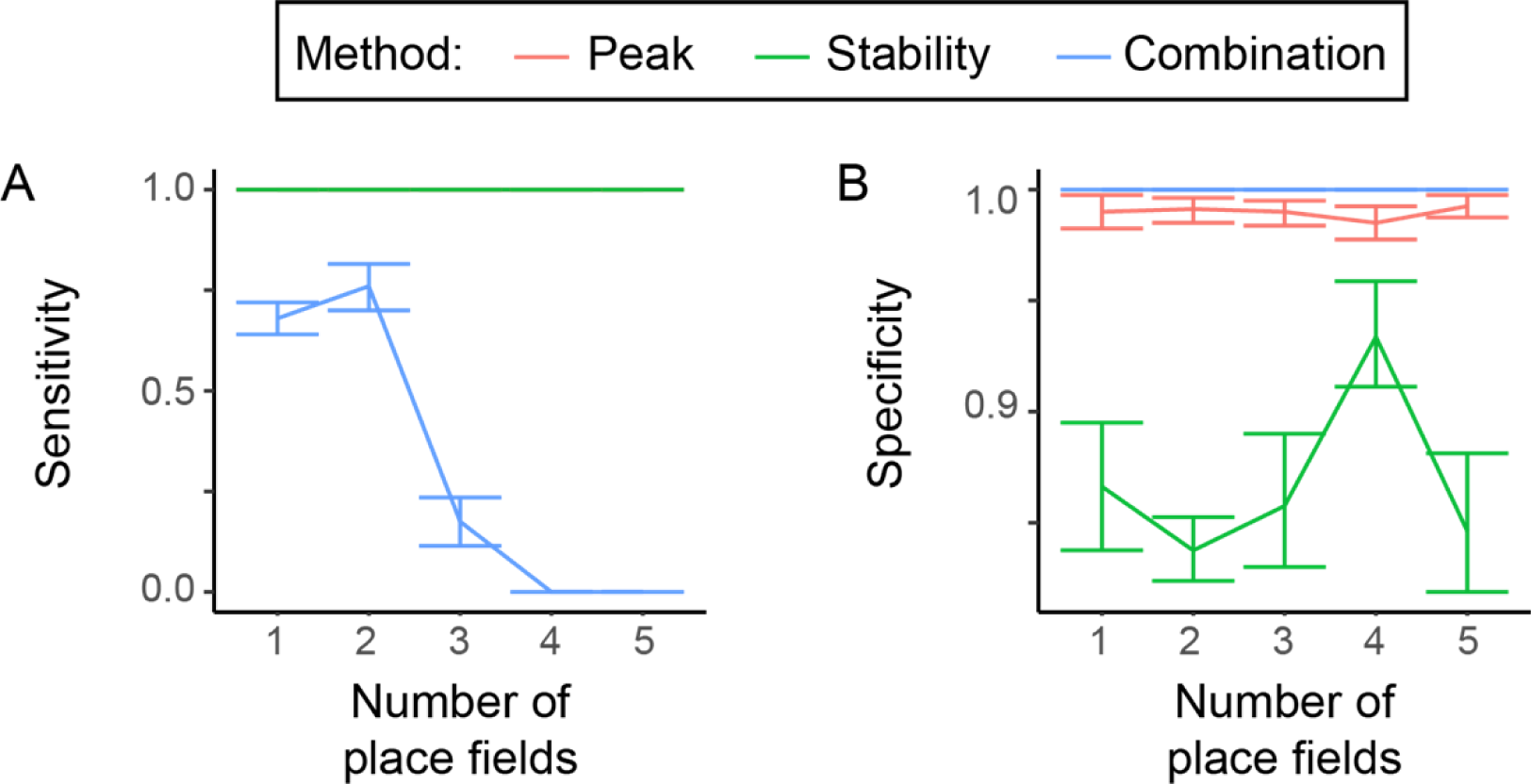
The effect of the number of place fields on detection. (A) Sensitivity and (B) specificity of methods as a function of the number of place fields. Data shown are the means of 10 trials with randomly created data using the set parameters, error bars show 95% confidence interval.

**S4 Fig.**
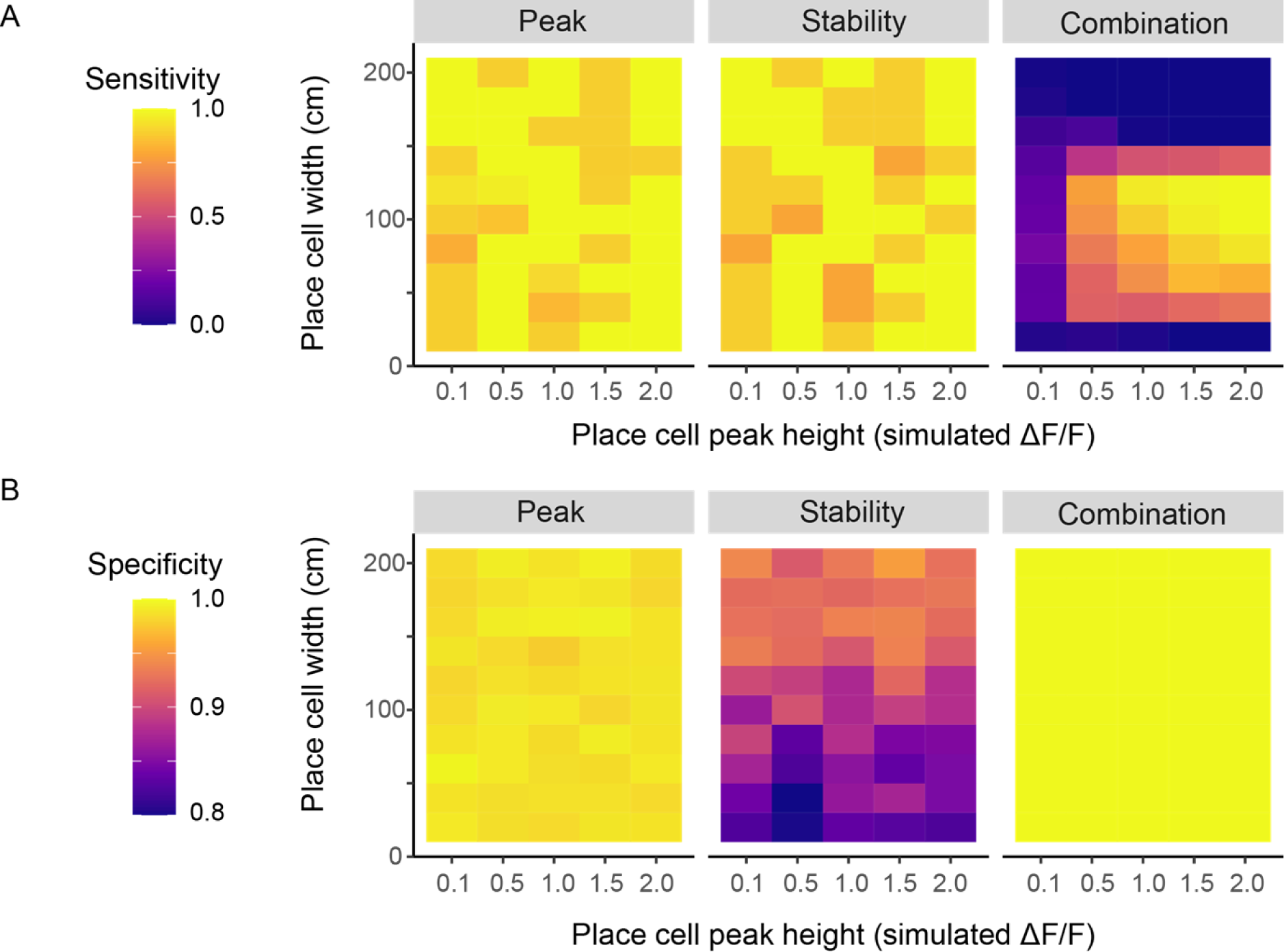
The combined effect place field width and peak. Sensitivity (A) and specificity (B) of the Peak, Stability, and Combination methods (as labelled) as a function of the peak of the place field and the width of the place field (up to 200 cm, the full environment length). Data shown are the means of 10 trials with randomly created data using the set parameters.

**S5 Fig.**
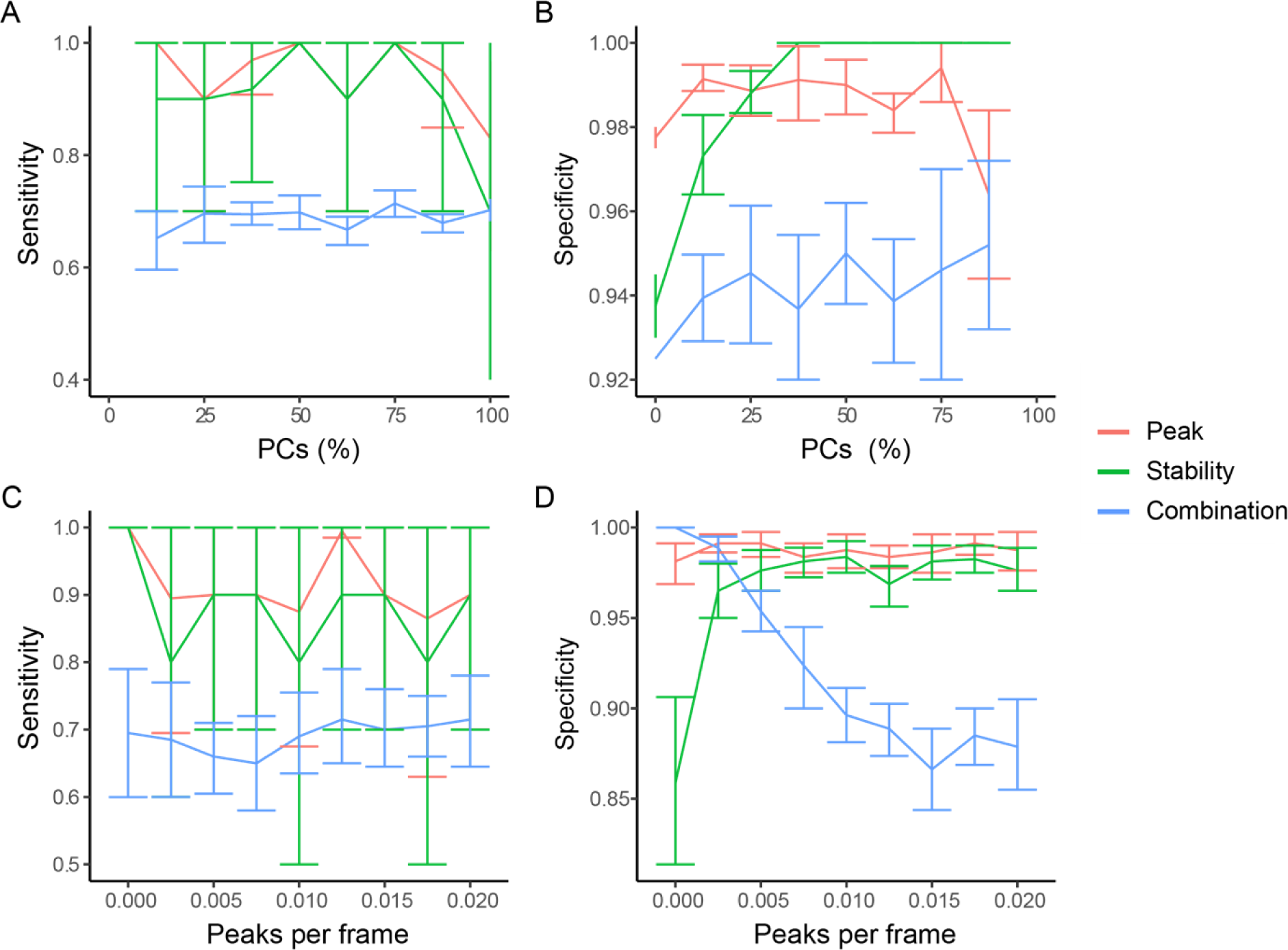
The effect of number of place cells and firing probability of non-place and place cells. Sensitivity (A) and specificity (B) of the Peak, Stability, and Combination methods as a function of the percentage of modelled place cells in the population. Altering the number of non-place cells increased specificity (i.e. fewer non-place cells, and therefore fewer total cells, were identified) when the number of place cells was increased beyond what is likely to be physiological (20-30%, [23]). Sensitivity (C) and specificity (D) of the Peak, Stability, and Combination methods as a function of the activity (peaks per frame) of the non-place cells. In the originally modelled datasets, the non-place cells all contained random background levels of Poisson noise with no firing, whereas the non-place cells in real data most likely do fire, though in a location-independent manner. We therefore introduced calcium peaks in the non-place cells with shapes and numbers of peaks modelled on our real data. There was no effect on sensitivity of increasing the activity of non-place cells, but the specificity of the Stability method increased and that of the Combination method decreased, with increasing non-place cell activity. These results cannot therefore explain why the Stability method er cells in our real data set. Data shown are the means of 10 trials with randomly created bars show 95% confidence intervals.

**S6 Fig.**
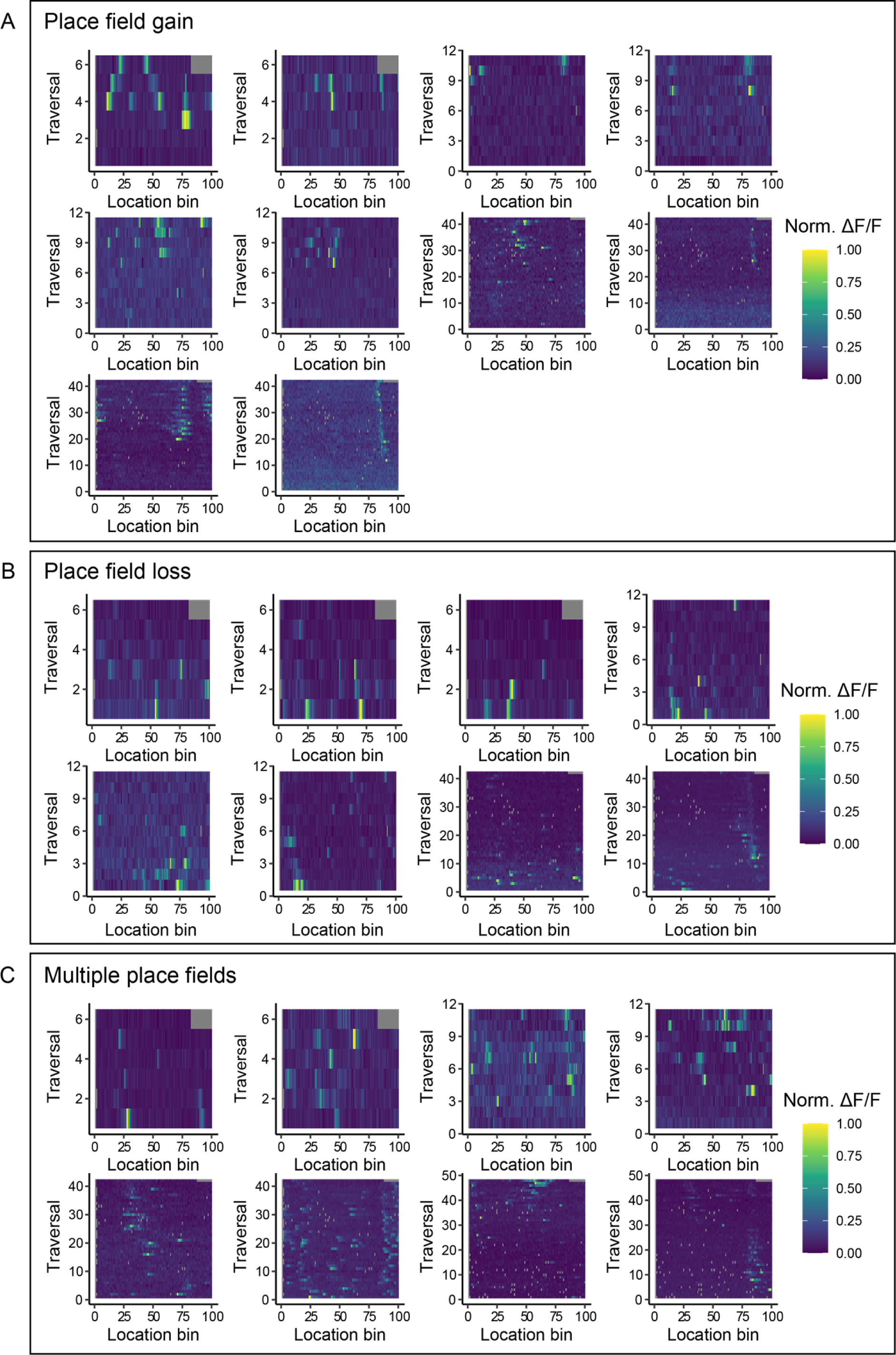
Examples of cells identified by the Peak method, but not the Stability method. Fluorescence ample cells that either (A) gain or (B) lose a place field, or (C) that have different place fferent times in the session. The colour represents the normalized fluorescence.

**S7 Fig.**
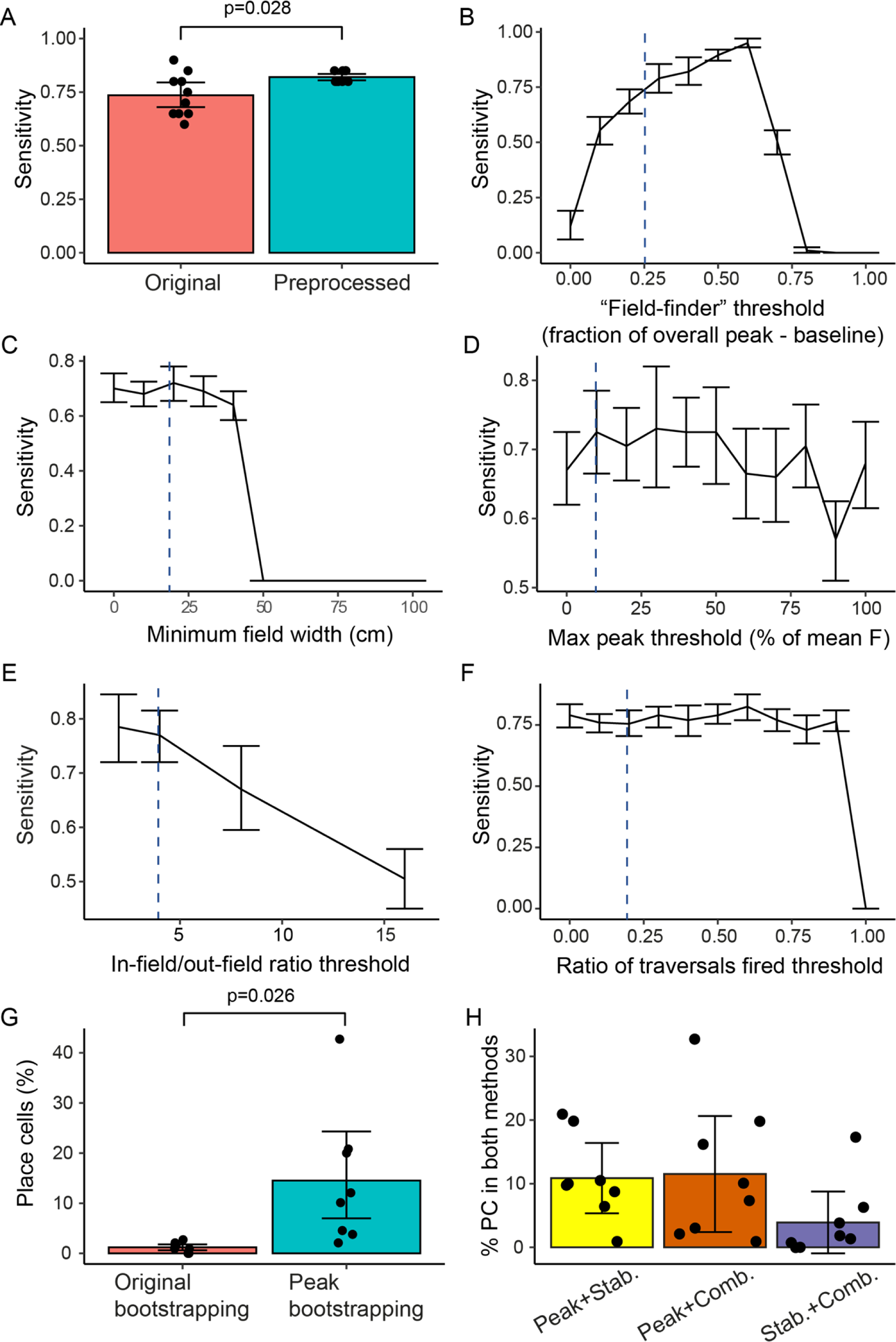
The effect of altering Combination method thresholds on identification of place cells in model and real data. (A) The sensitivity of the Combination method on detecting place cells in model data is increased when preprocessing is introduced, The sensitivity of the Combination method as a function of (B) in-field threshold (the minimum peak within a place field allowed as a fraction of the difference between the overall fluorescence peak in the data and the baseline), (C) minimum field width, (D) maximum fluorescence peak in the field, (E) in/out-field ratio, (F) ratio of traversals the cell fired in. (G) The percentage of ROIs in real data identified as place fields using Combination or Peak bootstrapping methods. (H) The percentage of ROIs identified as place cells by two methods, using the Peak bootstrapping method within the Combination method. Blue dashed lines indicate parameters used in published data and earlier analyses here. P-values are from linear mixed effect models, with the session included as a random effect.

